# A Two-Step Bioreactor for Lung Recellularization

**DOI:** 10.1101/2020.11.02.360792

**Authors:** Bethany Young, Leigh-Ann M. Antczak, Keerthana Shankar, Rebecca L. Heise

**Affiliations:** Department of Biomedical Engineering, Virginia Commonwealth University, 800 E. Leigh St, Room 1071, Richmond, VA 23219; Department of Physiology and Biophysics, Virginia Commonwealth University School of Medicine, 1101 East Marshall St, Richmond, Virginia 23298

**Keywords:** bioreactor, rotation seeding, lung tissue engineering, decellularized scaffold

## Abstract

Bioreactors for reseeding of decellularized lung scaffolds have evolved with a wide variety of advancements. These include biomimetic mechanical stimulation, constant nutrient flow, multi-output monitoring, and large mammal scaling. Although dynamic bioreactors are not new to the field of bioengineered lungs, ideal conditions during cell seeding have not been extensively studied or controlled. To address the lack of cell dispersal in traditional seeding methods, we have designed a two-step bioreactor. The first step rotates a seeded lung every 20 minutes at different angles to ensure 20 percent of cells are anchored to a particular location based on the known rate of attachment. The second step involves perfusion culture to ensure nutrient dispersion and cellular growth. Compared to statically seeded lungs followed by conventional perfusion, rotationally seeded lungs followed by perfusion had significantly increased dsDNA content and more uniform cellular distribution. This new bioreactor system will aid in recellularizing the lung and other geometrically complex organs for tissue engineering.

## Introduction

Designing methods for the culture of 3D tissue engineering requires strategies that do not typically apply to 2D culture. The intricate architecture necessary to create the natural tissue microenvironment in the lungs involves consideration of adequate cell seeding to all alveoli, airways, and capillaries, delivery of new nutrients, and tissue orientation during culture to ensure proper cell dispersal to all dimensions. Currently, there are several challenges hindering lung bioengineering from creating viable transplant alternatives, including insufficient scaffold cell coverage, partial cell differentiation, and lack of uniform organ function. Improving bioreactor designs and seeding techniques with the addition of cell-specific dynamic seeding protocols to current methods may aid in overcoming some of these challenges.

For the culture of tissue-engineered constructs, bioreactors improve several phases of regeneration: 1) cell seeding and dispersal, 2) delivery of fresh nutrients to enhance cell viability, and 3) applying a biomimetic mechanical stimulus to guide differentiation. Adequate simulation of healthy lung physiology depends on graft species origin and size. For rats specifically, *in vivo*, the maximum arterial flow rate would be between 10 and 20 mL/min and total lung capacity of 10-15 mL (1). The output parameters of designed bioreactors may not need to be as high as seen in healthy tissue but must resemble similar dynamic conditions. Design specifications for a lung bioreactor include a closed system to reduce contamination, ventilation with negative pressure to administer 0-15 breaths per minute, pulsatile perfusion at 4 mL/min, and constant oxygenation of the media (2). While ideal physiological conditions would ventilate the culturing lung with air through the trachea to achieve an air-liquid interface, during the initial stages of cell seeding, prematurely introducing air can cause epithelial detachment; therefore ventilation with media is advised for the first several weeks of culture (3).

Some of the most simple bioreactor systems are hollow fiber bioreactors that allow for basic perfusion of media or air around the structure for cell culture (4). More complex and controlled bioreactors are whole lung perfusion systems equipped with artificial silicone pleura and functional monitoring (2). Cell distribution onto and within scaffolds is improved by culturing in a dynamic bioreactor system to simulate the physiological environment and innovative protocols that allow for the most effective seeding of cells (5–7). Bioreactors have also been automated for both decellularization and recellularization procedures to ensure reproducibility. With many minor variations, the most common system performs cell seeding and culture with pulsatile or continuous vascular perfusion and negative pressure breathing (8,9).

For recellularization of acellular lung scaffolds, two main bioreactor designs have been developed to apply media flow and cell dispersal; a perfusion bioreactor system and a rotating wall vessel (RWV)(10,11). The most common device for cell seeding is the perfusion bioreactor system that seeds through static gravitational settling of a suspended cell solution or continuous flow of a cell suspension through the vasculature and airways. Another bioreactor designed to increase cell dispersal is the rotating wall vessel (RWV) bioreactor system (Figure 2.5C)(11). Within this system, scaffolds are perfused with cells and then centrifuged to disperse cells throughout the lung initially. Both of these designs have their limitations due to either cell settling during prolonged static culture periods or constant cell suspension flow, inhibiting the cells' ability to attach. Histological sections of recellularized lungs show the regional distribution and lack of complete alveolar or airway cell coverage. This indicates that cell seeding and dispersal techniques still need to be optimized during the first few hours.

The latest advances in both commercially available and custom lung bioreactor systems have extended the length of culture and reproducibility; however, with insufficient cell attachment and differentiation, current efforts have not resulted in fully functional tissue even after one month of culture (12,13). To accomplish bioengineered lung function, alternative seeding procedures need to be considered to combine with current bioreactor advancements (14). For these reasons, we have designed a bioreactor system that combines the beneficial aspects of all methods, with intermittent rotation tailored to cell attachment rates, a period of static culture to allow adhesions to mature, and finally perfusion and ventilation culture for mechanical conditioning.

The branching airway structures of the lungs are ideal for the efficient delivery of fluids and cells. Still, challenges arise with keeping cells stationary for attachment throughout all dimensions of all alveolar structures. Alveoli are spherical structures lined with a very thin alveolar epithelium that is vital to the function of the lung. The 200 µm structure is spherical and would require attachment or migration of approximately ten, 20 µm type 1 epithelial cells around the sphere (15). Due to the complex 3D structure, distribution of the cells within the alveoli and throughout all alveoli may require dynamic seeding regimens (11); however, during the first hours of culture, when cells are first attaching, periodically repositioning the lung may be needed for epithelial attachment. Given the importance of maximum attachment of delivered cells throughout recellularized lungs, the goal of this study is to increase cell dispersal using a rotational seeding method for *ex vivo* culture of decellularized lungs. Figure 1 gives an overview of the process.

**Figure 1.**
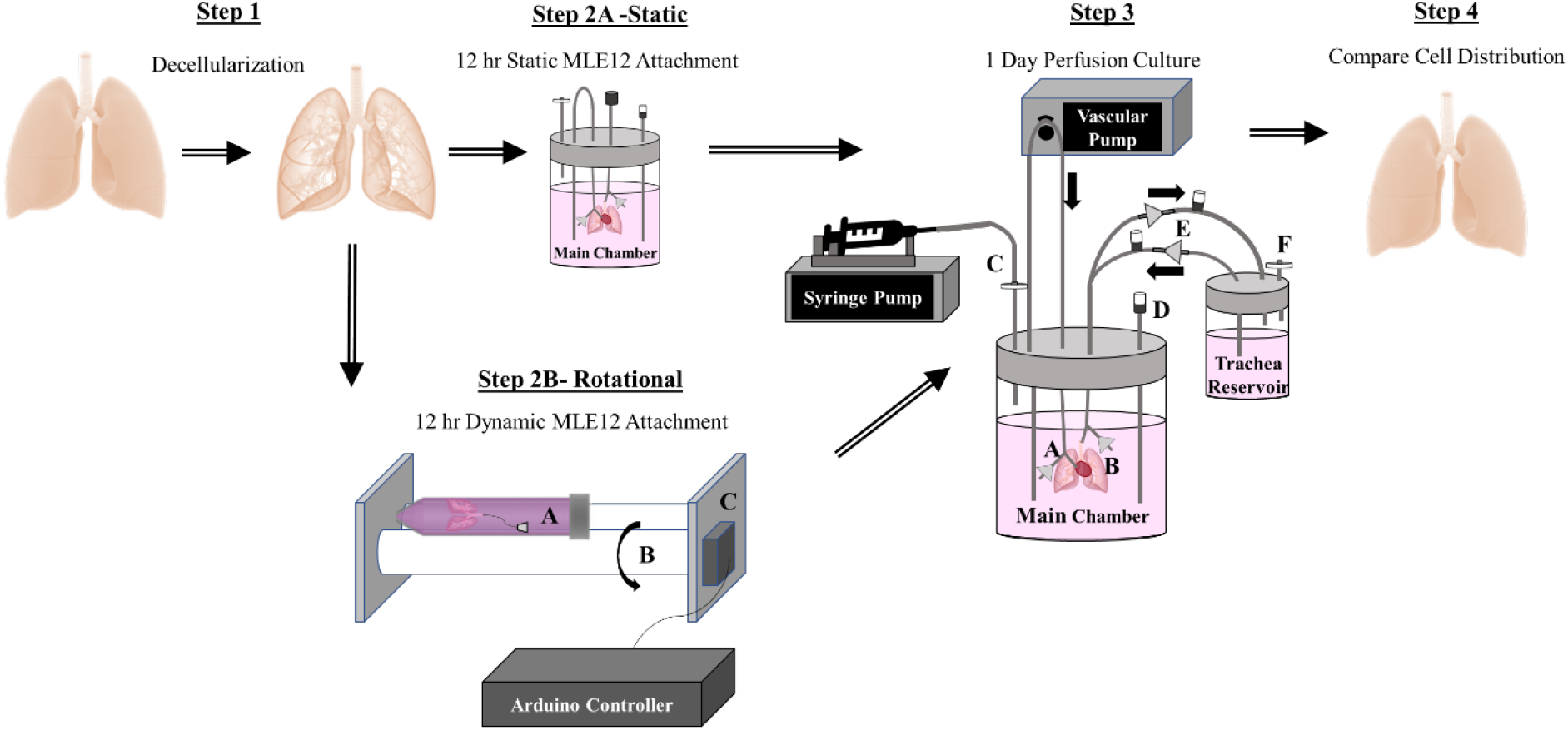
Overview of Cell Seeding and Lung Culture Methods. Rat lungs were first decellularized over a 3 day period. The resulting scaffolds were either statically or rotationally seeded for 12 hours. Statically-seeded lungs were transferred into the main chamber of the perfusion system and left at 37° C overnight. Alternatively, to encourage cell distribution, decellularized lungs were also placed in a 50 mL conical tube (Step 2B, part C) on two cylinders for periodic rotation. A stepper motor (Step 2B, part C) controlled by a programmed Arduino microcontroller rotated one cylinder (Step 2B, part B) and subsequently the lung within the tube systematically every 20 minutes.

## Materials and Methods

### Two-Step Bioreactor Design and Preparation

The first component of the two-step bioreactor is the rotational seeding system designed for improved attachment and dispersal of cells within decellularized lung scaffolds. The rotational system supports a 50 mL conical tube on two long cylinders (Figure 1). One cylinder was attached to a stepper motor programmed using an Arduino Mega controller system to rotate the tissue every 20 minutes for 3 hours. This rotation rate ensures uniform coverage of the airways over 3 hours using the cell coating routine illustrated in Figure 2. This rotation procedure allows attachment at every 45° angle of the lung throughout the entire culture. The degree of rotation between resting positions is large to allow for optimal cell movement throughout the tissue. After seeding completion, the lung was transferred to the second component of the two-step bioreactor system. The second component was a perfusion bioreactor system (Figure 1) for static overnight culture without vascular or airway perfusion. 24 hours after initial seeding, vascular perfusion through the pulmonary artery was initiated at 2 mL/minute by a pulsatile pump (CellMax). Media ventilation was also applied into the trachea at 1 breath/minute, driven by negative pressure within the main chamber, created by a syringe pump. One-way valves were placed into the breathing loop to allow new media exchange during ventilation. Before recellularization, the lung was rinsed 3 times with PBS with 5% Antibiotic/Antimycotic. All bioreactor components were autoclaved, rinsed with PBS with 5% Antibiotic/Antimycotic for 1 day, and handled within a sterile hood before use. 50 × 10^6^ MLE12s in 12 mL of media were perfused intratracheally. Inoculated lungs were either rotationally cultured for 3 hours in a 50 ml tube or placed directly into the perfusion bioreactor chamber for static culture overnight. After initial seeding on rotation bioreactor, as described in the bioreactor design, pulsatile vascular flow and media ventilation was administered for 24 hours.

**Figure 2.**
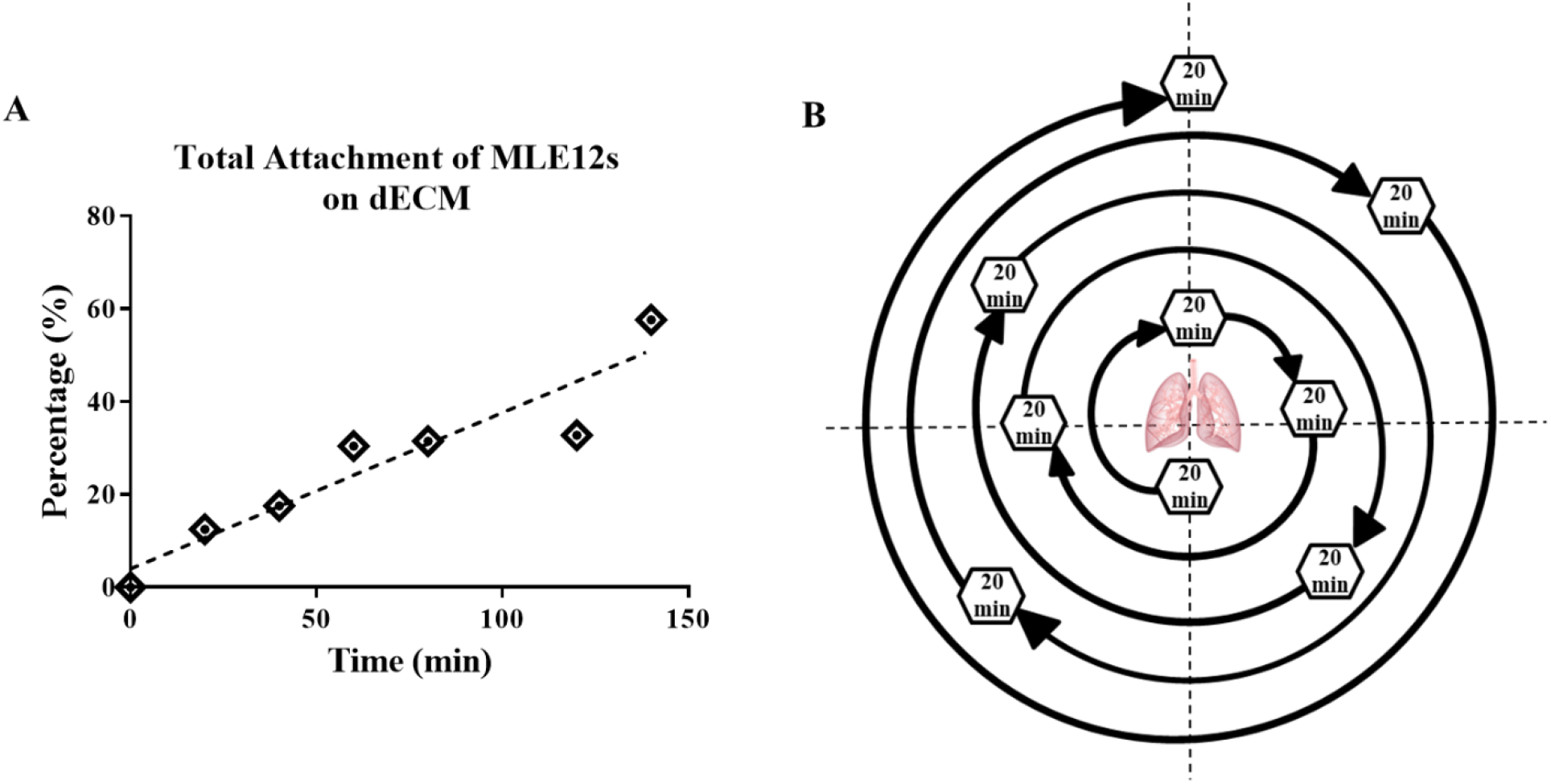
Rotational seeding movement pattern over 3 hours. The rate of MLE12 cell attachment to lung dECM coatings on TCP (A) shows ~15% of cells attach every 20 minutes. The rotation sequence (B) of the cell seeding bioreactor was programmed to rotate to the normal plane every 20-minutes. Large angles of rotation between each resting position allow for recoating the cells throughout the lung before reaching its 20-minute resting position. Data are presented as mean from 3 experimental replicates.

### Decellularization

Male and female Sprague Dawley rat lungs were decellularized by published methods (16). The trachea and pulmonary artery were cannulated to administer rinses and decellularization reagents. Each day over three days, the lung was perfused a minimum of three times with either 0.1% Triton X-100 (Fisher Scientific), 2% sodium deoxycholate (Sigma), and sodium chloride (Fisher Scientific), or a DNase solution and submerged overnight. After each overnight incubation and before perfusion of the next reagent, the lung was rinsed a minimum of three times with a PBS and penicillin/streptomycin solution.

### Cell Culture

Mouse alveolar type II cells (MLE12, ATCC) were cultured with HITES medium containing 2% fetal bovine serum at 37 °C with 5% CO_2_, according to the manufacturer’s protocols.

### MLE12 cell attachment rate assay

250,000 cells/cm^2^ were seeded onto 0.1 mg/mL porcine lung dECM coatings previously published by our laboratory (16–18). Every 20 minutes for 3 hours, unadhered cells and media were aspirated from the well and rinsed gently before fixation with 4 % PFA. Cell nuclei were stained using DAPI Prolong Gold Antifade Mounting agent, and the number of nuclei was counted within 3 pre-determined regions of each well using a Zeiss AxioObserver Z1 fluorescence microscope. Percent of cell attachment was calculated by dividing the number of cells attached by the number of cells seeded.

### Resazurin Reduction Assay

After 48 hours of perfusion culture, media was removed from the main chamber and replaced with 50 μM resazurin media solution. This solution remained within the lung for 4 hours until absorbance (544 nm (ex)/590 nm (em)) levels of the resazurin solution and qualitative images were taken to evaluate metabolic activity of the cells. Absorbance values were subtracted from the non-metabolized stock concentration of the 50 μM resazurin solution.

### Histology Imaging and Quantification

Each lung (right or left) was divided into anterior or posterior portions, that were further divided into upper and lower regions. Dissected tissue for histology and other assays were chosen based on resazurin cell viability throughout the lung, to attempt to isolate regions with similar cell viability for histology and Picogreen. An entire anterior or posterior portion of each lung was fixed with 4% paraformaldehyde, embedded, and sectioned through the frontal plane to see the distribution of cells throughout the entirety of the lung. Two histological sections, several millimeters apart were taken from each tissue region for staining with hematoxylin and eosin (H & E) and imaged using a Vectra Polaris Automated Quantitative Pathology Imaging System.

20x images from five set regions (top, bottom, right, left, and center) of one histological section were randomly selected and analyzed using Image J. A mask of each image was created with a set threshold range from 50 to 150. This created a binary image containing only the cell nuclei, which was verified by hand counting. After separating touching nuclei and removing any background signal smaller than 15 pixels, the analyzed particles function was used to count the cell number.

### dsDNA quantification

A Picogreen dsDNA quantification assay (ThermoFisher) was performed on the remaining portions of the lungs not used for histological sectioning. Each sample was normalized to its wet weight and graphed as total dsDNA throughout the tissue.

### Statistical Analysis

All data are presented as mean +/− standard deviation with an N ≥ 3 unless otherwise stated. Statistical significance was determined by a Two or One-way ANOVA and a Tukey’s multiple comparisons tests using GraphPad Prism.

## 5.3 Results

### Bioreactor design

To enhance cell dispersal and mimic the mechanics, temperature, and pH within the lungs, we have designed a novel bioreactor system. The first portion of the bioreactor system is a rotational seeding unit (Figure 1) that will turn the lung every 20 minutes in a pattern illustrated in figure 5.1B for 3 hours during the initial cell attachment period. Large degrees of rotation was used between resting periods to tumble the cells through multiple regions of the lungs before settling at the desired location. The duration of the resting periods was determined by the rate of MLE12 attachment (Figure 2) to allow 10 percent of cells to attach at each position. This aided in the attachment of cells more evenly throughout the lung. Seeding efficiency was characterized by H & E staining and Picogreen dsDNA quantification. After 3 hours of cell attachment, the lung was moved to the perfusion bioreactor (Figure 1) for mechanical conditioning and media perfusion. The perfusion bioreactor design was adapted from a previously published design(8,19) without the endothelial seeding chamber.

### Resazurin and Picogreen assessment of cell distribution within cultured lungs

50 million MLE12 cells were cultured into decellularized rat lungs with rotational or static seeding to assess if periods of dynamic culture can increase cell dispersal throughout the lung. Resazurin was perfused through the airways and cultured for 4 hours to allow for metabolites to gradually change the color of the resazurin solution from a dark blue to pink for whole-lung cell viability assessments. After 48 hours of culture, rotationally seeded lungs exhibited a more uniform pink or light purple color distribution, correlating to viable cells throughout, while static seeding causes several distinct regions with either viable cells (pink) or dark purple to blue (less cellular metabolism, Figure 3. Resazurin absorbance of the lung perfusate (Figure 3) was collected from the tracheal cannula and showed increased cell viability with rotational seeding. There was no significant increase between rotationally and statically seeded lungs because the collection of the perfusate from the trachea had mixed with the resazurin from all regions of the lung upon removal. To better understand where cells attached with both seeding methods, after 48 hours of culture, the lung was dissected into several regions for assessment. Summing the dsDNA within all areas of each lung showed total dsDNA concentrations (Figure 3) that resembled the results seen by resazurin absorbance, as expected.

**Figure 3.**
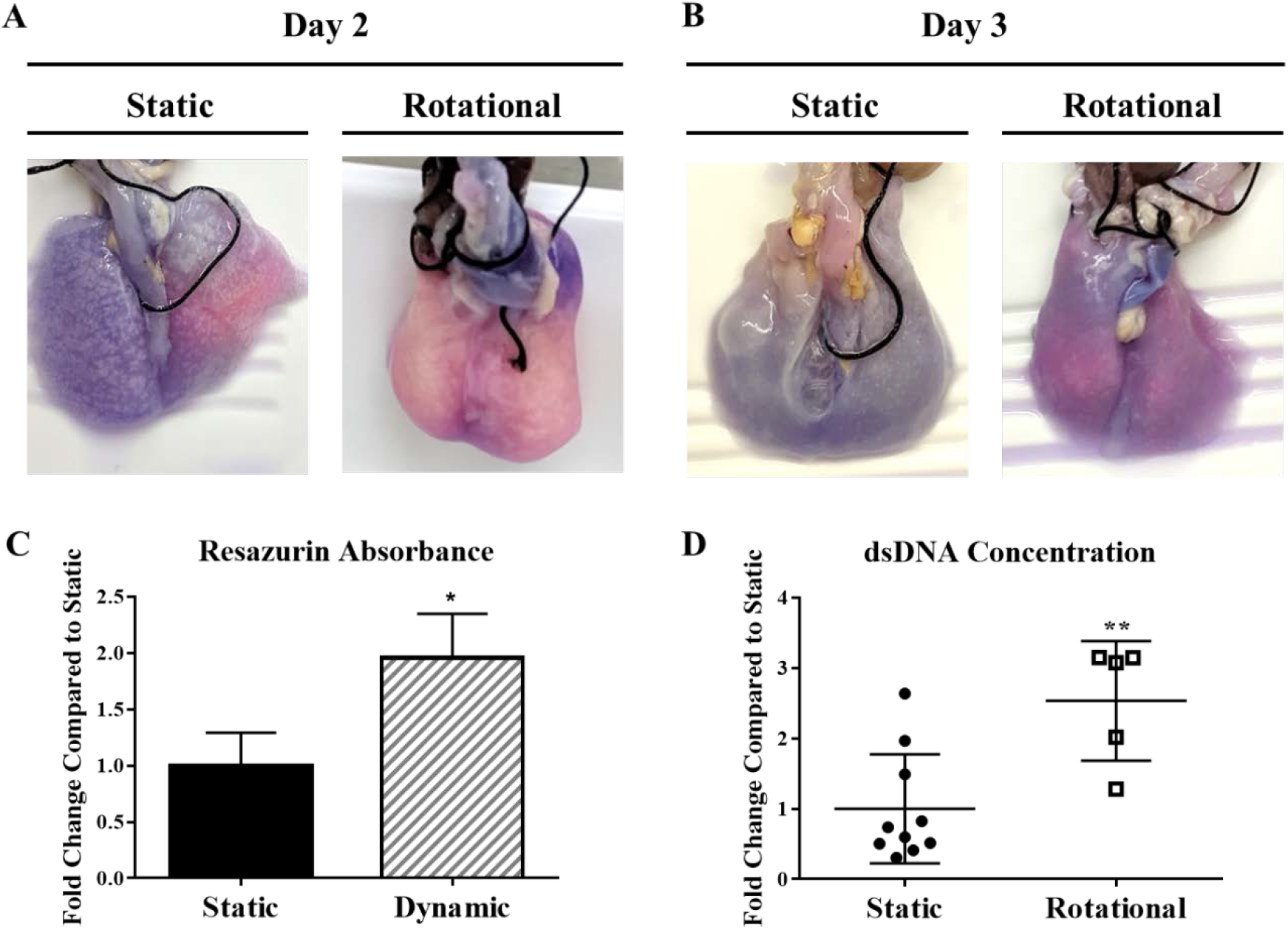
Whole Lung Cellular Distribution. Images of reseeded lungs after resazurin perfusion at day 2 (A) and day 3 (B). Pink regions show areas with more cellular activity and dark blue/purple areas with less cells. Quantification of resazurin absorbance from each set of lungs after 3 days of culture (C). dsDNA quantification after 3 Days (D) compares the concentration distribution between static to rotationally seeded lungs with each circle or square representing a concentration from either an anterior or posterior portion of each lung. All lungs but one had more than two regions within one lung represented within the data. Data are presented as mean +/− standard deviation from 3-4 reseeded lungs. *, ** indicates p < 0.05, p< 0.01 respectively.

### Cell dispersal within dynamically culture lungs

Histology was also done to verify the findings from resazurin and Picogreen cell distribution studies. After 48 hours of culture (Figure 4), there were increases in cell attachment and distribution. Cell morphology within both conditions does not show cell spreading with some clumping of cells within the airways, which may be attenuated with longer culture times.

**Figure 4.**
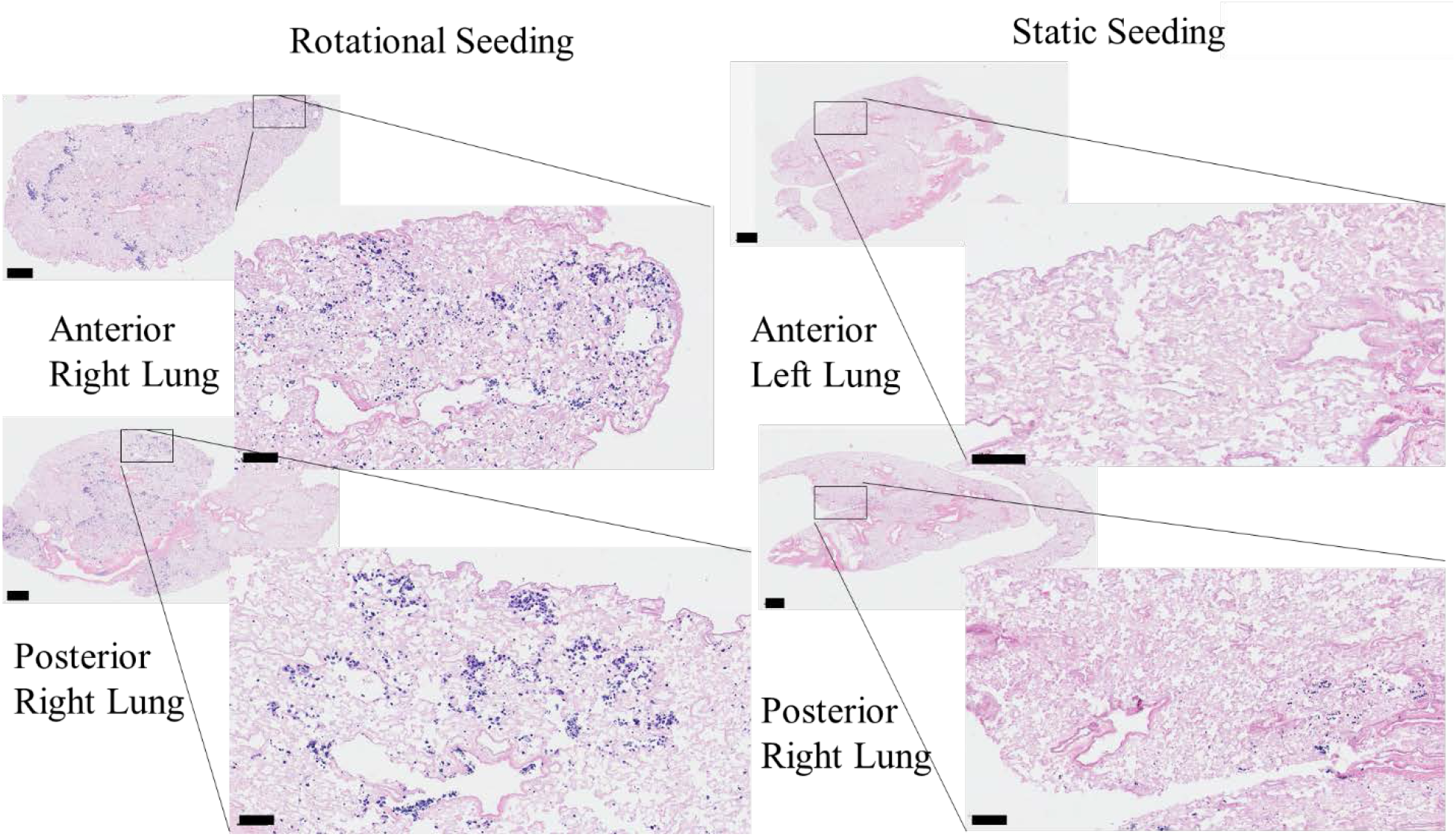
Histological analysis of cell attachment after static or dynamic seeding. Representative images from the histological sections from either the anterior or posterior regions of the left or right lungs are shown. Scale bars on larger images indicate 100 μm, and scale bars on smaller, magnified images indicate 600 μm.

## 5.4 Discussion

**S**everal other groups have concluded that dynamic seeding of cells onto biomaterials is necessary to enhance whole lung bioengineering; however, implementation of these methods has faced unanswered challenges. The perfusion bioreactor system is done without flow for the first 12 hours to 2 days to allow for seeding. With this method, gravity and cell settling can drastically affect cell dispersal. Within the same system, cellular seeding has also been done in the presence of either vascular or airway flow, continuously moving cells and media. This method would increase the dispersal of cells; however, suspension cultures can disrupt cell attachment rates (20,21). The RWV system initially disperses cells throughout the lung and then statically cultures the seeded tissue for 1 to 4 days before initiating rotational culture (11,22,23). This system mainly increases nutrient flow and creates microgravity throughout the culture; however, since the lungs are statically cultured before horizontal rotation, the coating of cells throughout the lung is limited by the sedimentation and regional seeding of cells. Even if the rotation began during the time of seeding, constant rotation at 20 RPM would not allow cells to remain stationary for long enough for cells to adhere to the matrix. These systems have been designed and commercialized for both rodent lung culture and more complex porcine and human-sized organs (2,8,20). Advances in both commercially available and custom lung bioreactor systems include artificial pleural membranes, gas exchange units, waste removal, real-time monitoring of oxygen and lung mechanics, and high-throughput automation (12). These advancements have extended the length of culture and reproducibility; however, with insufficient cell attachment and differentiation, functional tissue formation has yet to be achieved.

*In vivo*, migratory cells such as immune or metastatic cells require flow to initiate cell adhesion. In the case of epithelial cells, laminar flow can decrease the rate of attachment (21) and even increase metastatic potential (24). Alternatively, to ensure sufficient epithelial attachment, periods of static culture are required to allow for cell adhesion complexes to be formed. Cells first make attachments to their ECM substrates through integrin-based focal adhesions. Epithelial cells then spread, reorganize their cytoskeleton, and initiate more integrin interactions with the ECM to strengthen adhesion over time. Adhesion complexes begin cooperatively binding after just one minute, but it takes up to 10 to 20 minutes after seeding for the cell adhesion structures to withstand substantial shear forces (25). Once epithelial attachments have matured, fluid flow and shear stresses become an essential driver of cell differentiation. After attachment, a long-term culture within a perfusion bioreactor system is used to distribute nutrients and apply biophysical stimuli. While perfusion and RWV systems provide beneficial advanced media dispersal and mechanical cues that aid in tissue maturation they require refining concerning seeding procedures. The bioreactor design described in this study consolidates the innovative properties of these traditional systems while overcoming their limitations in cell dispersal and attachment.

In this study, we have developed an increased dispersal cell seeding protocol that overcomes the limitations of traditional bioreactor methods to be used in tandem with other long-term culture techniques. This two-phase culture system includes a rotational cell attachment step to increase cell distribution throughout the lung, followed by traditional *ex vivo* lung perfusion culture. Over 60 different cell types can be found in the lungs, excluding the circulatory cells. These cell types represent a diversity of functions and therefore have varying attachment and migration characteristics (26). Different requirements for cell adhesion are needed and are dependent on the cell's specific applications. Alterations to this seeding method, such as adjustments to the elapsed time between rotations as well as the speed of the rotation, can create custom seeding routines based on the differing cell attachment characteristics.

Resazurin metabolic assays have been used as an efficient, non-invasive way to assess tissue viability during *ex vivo* culture (25,27,28). With perfusion into the seeded scaffold, cells will encounter resazurin and undergo reduction into a pink colored resorufin. Pink and purple regions with high cellular viability will easily be distinguished from dark blue regions with little cell content. Rotationally seeded lungs exhibited pink and light purple regions throughout the lung scaffold. In contrast, statically seeded scaffolds had some bright pink areas that rapidly transitioned to a dark purple or blue color. The average resazurin absorbance showed similar cell viability results between the two groups. Picogreen dsDNA concentration differences between static and rotationally seeded lungs showed that rotational seeding was statistically more significant than static seeding.

To confirm our previous findings, histology through the frontal plane of the cultured lung shows a broader distribution of cells throughout dynamically seeded lungs. Cells within statically seeded lungs exhibit sedimentation in a small portion of the lung, limiting overall function after long-term culture. Cell clumping within alveolar and airway spaces and rounded cell morphologies may indicate that longer culture times are needed. Various epithelial cell lineages or stem cell populations may also be used in the future to induce phenotypic morphologies and increase airway and alveoli coverage.

The purpose of this research is to develop a bioreactor system that can be used with all future tissue engineering advancements and propel future technologies by starting with an efficient seeding mechanism. Since cellular distribution and attachment were the main goal of these studies, there are some limitations of the study. This study only shows short term cell distribution over 3 days and, therefore, does not provide context for cell migration, proliferation, or differentiation. Also, the nature of a two-step bioreactor causes challenges concerning sterility. Within this bioreactor design, the tissue must be transferred from the rotational seeding container to the conventional bioreactor chamber after the seeding period is complete. For that reason, two rotationally seeded lungs experienced contaminations. Future studies should incorporate a design modification to create a fully closed system for the rotational seeding portion and the transfer to the conventional bioreactor. These studies can also include other upgrades and techniques for long-term tissue culture to study more complex tissue transplantation challenges such as cellular differentiation, barrier integrity, longer-term function.

## Conclusions

Careful consideration of epithelial attachment *in vitro* was used to determine the ideal attachment rate within a whole lung scaffold and improved cell coverage. These studies help to inform all types of 3D culture methods and emphasize the importance of intermittent motion in cellular seeding. The development of a portable, rotation system that is not in direct contact with the lung or media allows for reduced contamination and adaptation of these methods to biomaterial applications. Overall, these are promising results highlighting the importance of dynamic culture in *ex vivo* cell culture to increase recellularization efficiency.

## References

1. Gilpin SE, Charest JM, Ren X, Ott HC. Bioengineering Lungs for Transplantation. Thorac Surg Clin. 2016 May 1;26(2):163–71.

2. Raredon MSB, Rocco KA, Gheorghe CP, Sivarapatna A, Ghaedi M, Balestrini JL, et al. Biomimetic Culture Reactor for Whole-Lung Engineering. BioResearch Open Access. 2016 Apr 11;5(1):72–83.

3. Petersen TH, Calle EA, Colehour MB, Niklason LE. Bioreactor for the Long-Term Culture of Lung Tissue. Cell Transplant. 2011 Aug 1;20(7):1117–26.

4. Guyot A, Hanrahan JW. ATP release from human airway epithelial cells studied using a capillary cell culture system. J Physiol. 2002 Nov 15;545(Pt 1):199–206.

5. Gilpin SE, Li Q, Evangelista-Leite D, Ren X, Reinhardt DP, Frey BL, et al. Fibrillin-2 and Tenascin-C bridge the age gap in lung epithelial regeneration. Biomaterials. 2017 Sep 1;140:212–9.

6. Le AV, Hatachi G, Beloiartsev A, Ghaedi M, Engler AJ, Baevova P, et al. Efficient and Functional Endothelial Repopulation of Whole Lung Organ Scaffolds. ACS Biomater Sci Eng. 2017 Sep 11;3(9):2000–10.

7. Ren X, Moser PT, Gilpin SE, Okamoto T, Wu T, Tapias LF, et al. Engineering pulmonary vasculature in decellularized rat and human lungs. Nat Biotechnol. 2015 Oct;33(10):1097–102.

8. Bonvillain RW, Scarritt ME, Pashos NC, Mayeux JP, Meshberger CL, Betancourt AM, et al. Nonhuman Primate Lung Decellularization and Recellularization Using a Specialized Large-organ Bioreactor. JoVE J Vis Exp. 2013 Dec 15;(82):e50825–e50825.

9. Ott HC, Clippinger B, Conrad C, Schuetz C, Pomerantseva I, Ikonomou L, et al. Regeneration and orthotopic transplantation of a bioartificial lung. Nat Med. 2010 Aug;16(8):927.

10. Petersen TH, Calle EA, Zhao L, Lee EJ, Gui L, Raredon MB, et al. Tissue-Engineered Lungs for in Vivo Implantation. Science. 2010 Jul 30;329(5991):538–41.

11. Crabbé A, Liu Y, Sarker SF, Bonenfant NR, Barrila J, Borg ZD, et al. Recellularization of Decellularized Lung Scaffolds Is Enhanced by Dynamic Suspension Culture. PLOS ONE. 2015 May 11;10(5):e0126846.

12. Charest JM, Okamoto T, Kitano K, Yasuda A, Gilpin SE, Mathisen DJ, et al. Design and validation of a clinical-scale bioreactor for long-term isolated lung culture. Biomaterials. 2015 Jun;52:79–87.

13. Nichols JE, Francesca SL, Niles JA, Vega SP, Argueta LB, Frank L, et al. Production and transplantation of bioengineered lung into a large-animal model. Sci Transl Med. 2018 Aug 1;10(452):eaao3926.

14. Joan E Nichols, Stephanie P Vega, Lissenya B Argueta, Jean A Niles. Modeling the lung: Design and development of tissue engineered macro- and micro-physiologic lung models for research use.

15. Wikswo JP, Curtis EL, Eagleton ZE, Evans BC, Kole A, Hofmeister LH, et al. Scaling and systems biology for integrating multiple organs-on-a-chip. Lab Chip. 2013 Aug 13;13(18):3496–511.

16. Pouliot RA, Link PA, Mikhaiel NS, Schneck MB, Valentine MS, Kamga Gninzeko FJ, et al. Development and characterization of a naturally derived lung extracellular matrix hydrogel. J Biomed Mater Res A. 2016 Mar 25;

17. Link PA, Pouliot RA, Mikhaiel NS, Young BM, Heise RL. Tunable Hydrogels from Pulmonary Extracellular Matrix for 3D Cell Culture. JoVE J Vis Exp. 2017 Jan 17;(119):e55094–e55094.

18. Young BM, Pouliot RA, Link PA, Park HE, Kahn AR, Shankar K, et al. Porcine Lung-Derived Extracellular Matrix Hydrogel Properties Are Dependent on Pepsin Digestion Time. Tissue Eng Part C Methods. 2020 Jun;26(6):332–46.

19. Calle EA, Petersen TH, Niklason LE. Procedure for Lung Engineering. J Vis Exp JoVE [Internet]. 2011 Mar 8 [cited 2017 May 17];(49). Available from: http://www.ncbi.nlm.nih.gov/pmc/articles/PMC3197323/

20. Nichols JE, La Francesca S, Vega SP, Niles JA, Argueta LB, Riddle M, et al. Giving new life to old lungs: methods to produce and assess whole human paediatric bioengineered lungs. J Tissue Eng Regen Med. 2017 Jul 1;11(7):2136–52.

21. Karuri NW, Liliensiek S, Teixeira AI, Abrams G, Campbell S, Nealey PF, et al. Biological length scale topography enhances cell-substratum adhesion of human corneal epithelial cells. J Cell Sci. 2004 Jul 1;117(15):3153–64.

22. Cortiella J, Niles J, Cantu A, Brettler A, Pham A, Vargas G, et al. Influence of Acellular Natural Lung Matrix on Murine Embryonic Stem Cell Differentiation and Tissue Formation. Tissue Eng Part A. 2010 Apr 19;16(8):2565–80.

23. Culturing and Applications of Rotating Wall Vessel Bioreactor Derived 3D Epithelial Cell Models [Internet]. [cited 2019 Apr 14]. Available from: https://www.ncbi.nlm.nih.gov/pmc/articles/PMC3567125/

24. Rizvi I, Gurkan UA, Tasoglu S, Alagic N, Celli JP, Mensah LB, et al. flow induces epithelial-mesenchymal transition, cellular heterogeneity and biomarker modulation in 3D ovarian cancer nodules. Proc Natl Acad Sci. 2013 May 28;110(22):E1974–83.

25. Taubenberger A, Cisneros DA, Friedrichs J, Puech P-H, Muller DJ, Franz CM. Revealing Early Steps of α2β1 Integrin-mediated Adhesion to Collagen Type I by Using Single-Cell Force Spectroscopy. Mol Biol Cell. 2007 Feb 21;18(5):1634–44.

26. Price AP, England KA, Matson AM, Blazar BR, Panoskaltsis-Mortari A. Development of a Decellularized Lung Bioreactor System for Bioengineering the Lung: The Matrix Reloaded. Tissue Eng Part A. 2010;16(8):2581–91.

27. Uzarski JS, DiVito MD, Wertheim JA, Miller WM. Essential Design Considerations for the Resazurin Reduction Assay to Noninvasively Quantify Cell Expansion within Perfused Extracellular Matrix Scaffolds. Biomaterials. 2017 Jun;129:163–75.

28. Gilpin SE, Charest JM, Ren X, Tapias LF, Wu T, Evangelista-Leite D, et al. Regenerative potential of human airway stem cells in lung epithelial engineering. Biomaterials. 2016 Nov 1;108:111–9.

